# Bronchial epithelial IB3-1 cells exposed to the anti-SARS-CoV2 BNT162b2 vaccine: induction of apoptosis is partially reversed by Aged Garlic Extract (AGE) and its bioactive components S-allyl-Cysteine and S1-propenyl-l-Cysteine

**DOI:** 10.64898/2026.09.19.752820

**Authors:** Federica di Padua, Giovanni Marzaro, Enzo Agostinelli, Roberto Gambari, Alessia Finotti

## Abstract

The Spike protein (S-protein) of the Severe Acute Respiratory Syndrome Coronavirus-2 (SARS-CoV2) exhibits important and negative effects on human cells. Accordingly, the adverse effects of COVID-19 mRNA vaccines could be due to the presence of circulating spike and need to be mitigated. One of the biological activities of the Spike protein (and, indirectly, of the Spike-producing vaccines) is the high induction of apoptosis and pro-inflammatory gene expression in different cellular systems. We have recently demonstrated that Aged Garlic Extract (AGE) and its components S-allyl-Cysteine (SAC) and S1-propenyl-I-Cysteine (S1PC) are potent inhibitors of the expression of pro-inflammatory genes induced by treatment of the bronchial epithelial IB3-1 cells with the Spike RNA-based BNT162b2 vaccine. The main purpose of the present study was to determine (a) whether treatment of the IB3-1 bronchial epithelia cells with the BNT162b2 vaccine is associated with activation of the apoptotic pathway and (b) whether AGE, SAC and S1PC have any effect of therapeutic relevance on this experimental model system. We have exposed IB3-1 cells to increasing amounts of the BNT162b2 vaccine and, after 3 days of culture, apoptosis was assessed using the Annex V and the Caspase 3/7 apoptosis assays. The results obtained demonstrated highly significant increase of the percentage of apoptotic cells after treatment of IB3-1 cells with the BNT162b2 vaccine. The percentage of apoptotic cells was found to be significantly reduced in cell populations treated with the BNT162b2 vaccine in the presence of AGE, SAC and S1PC. The main conclusion of the results of our study is that Aged Garlic Extracts, and its components S-allyl-Cysteine and S1-propenyl-l-Cysteine are able to mitigate the pro-apoptotic effects of the Spike-producing BNT162b2 vaccine using the bronchial epithelial IB3-1 cell line as experimental model system. The efficacy of the reversion of BNT162b2 induced apoptosis suggests that AGE, SAC and S1PC should be considered for the development of protocols aimed at counteracting apoptosis induced by the Spike mRNA-based COVID-19 vaccines. This might be of potential clinical relevance, since one of the biological adverse effects of Spike protein (and, indirectly, of the Spike-producing vaccines) is the high induction of apoptosis in different cellular systems.

## INTRODUCTION

The Spike protein (S-protein) of the Severe Acute Respiratory Syndrome Coronavirus-2 (SARS-CoV2) plays a very important role in the pathogenesis of Coronavirus Disease-19 (COVID-19) (1). Accordingly, several research efforts have been conducted to determine biological activities of the S-protein. In this context, there is general agreement on the fact that the SARS-CoV-2 Spike protein has important effects on human cells (1–3). This has been reviewed and appropriately discussed by Trougakos et al. (1), who hypothesized that adverse effects of COVID-19 mRNA vaccines could be due to the presence of circulating Spike. The incidence, clinical presentation, risk factors, diagnosis, and management of different adverse effects and their possible mechanisms have been discussed by Gopalaswamy et al., who concluded that the potential ambivalence of human response post-COVID-19 vaccination necessitates the need to mitigate the adverse side effects (3). In this respect, Meo et al. compared the biological, pharmacological characteristics and adverse effects of Pfizer/BioNTech and Moderna Vaccines. The expected conclusion was that the induction of anti-SARS-CoV-2 antibodies is associated with the production of high levels of the Spike protein (S-protein) (4).

In respect to possible adverse effects, one of the biological activities of the Spike protein (and, indirectly, of the Spike-producing vaccines) is the high induction of apoptosis in different cellular systems (5–8). For instance, Barhoumi et al. found that the Spike protein induces apoptosis in THP-1-Like-Macrophages (6). In addition, Yamamoto et al. reported that the SARS-CoV-2 spike protein promotes taste cell apoptosis and the release of the apoptosis-related cytokine TNF-α, implicating its contribution to the taste malfunction caused by COVID-19 (7). These data support the concept that the induction of apoptosis by the SARS-CoV-2 Spike protein is usually associated with deep alteration(s) of cellular functions. For instance, Wu et al. found that SARS-CoV-2 infects human pancreatic β cells in COVID-19 patients and selectively infects human islet β cells in vitro. This study demonstrated that SARS-CoV-2 infection attenuates pancreatic insulin levels and secretion and induces β cell apoptosis (8). In general, similar effects were found in SARS-CoV-2 infected cells, as well as in cells treated with recombinant Spike or Spike producing COVID-19 RNA-based vaccine such as the Pfizer BNT162b2 (9) and the Moderna mRNA-1273 (10) vaccines.

Notably, the Spike-mediated induction of cellular apoptosis is particularly relevant in the case of the effects of Spike or Spike-producing vaccines on T-cells. In this respect, Gimenez et al. found that monocytic reactive oxygen species-induced T-cell apoptosis impairs cellular immune response to SARS-CoV-2 mRNA vaccine (11). Accordingly, André et al. were able to demonstrate that low quantity and quality of anti-spike humoral response was linked to CD4 T-cell apoptosis in COVID-19 patients (12). Therefore, the mitigation of these adverse side effects is highly needed in order to maximize the beneficial effects of Spike producing RNA-based vaccines on COVID-19 patients.

In this respect, Roy et al. reported that *Eupatorium perfoliatum* prevents and alleviates SARS-CoV-2 Spike protein-induced lung inflammatory response and apoptosis (13). In this study, the treatment with S protein resulted in disruption of lung epithelial cell morphology, activation of the inflammasome, increased cell apoptosis, elevated pro-inflammatory cytokines, and extracellular matrix remodeling. Roy et al. explored the therapeutic potential of *Eupatorium perfoliatum* and found that it was able to mitigate most of the Spike-induced alterations, such as hyperinflammation and apoptosis, suggesting an interplay between hyperinflammation and apoptosis (13).

We have recently demonstrated that Aged Garlic Extract (AGE) and its components S-allyl-Cysteine (SAC) and S1-propenyl-l-Cysteine (S1PC) are potent inhibitors of the expression of pro-inflammatory genes induced by treatment of the bronchial epithelial cells with the Spike RNA-based BNT162b2 vaccine (14,15).

Among the possible biochemical targets of SAC and S1PC, Gasparello et al. (14) and Papi et al. (15) have reported molecular docking and molecular dynamics data supporting the concept that SAC and S1PC bind to Toll-like receptor-4 (TLR4), thereby interfering with the TLR4/NFkB signaling pathway (14,15). Notably, the SARS-CoV-2 spike protein binds to Toll-like receptor 4 (TLR4), triggering downstream MyD88-dependent NF-κB activation (16,17). This signaling cascade drives severe cellular hyperinflammation, oxidative stress, and subsequent apoptosis (16–18).

Among the several experimental model systems useful to detect activity of natural compounds on Spike-induced effects, we have extensively studied human bronchial epithelial IB3-1 cells treated with the SARS-CoV2-Spike or Spike-producing BNT162b2 vaccine (14,15,19,20). IB3-1 cells have been derived from a patient with cystic fibrosis carrying the *CFTR* ΔF508/W1282X genotype. The cells were immortalized by transfection with an adenovirus 12/SV40 hybrid virus (21), resulting in a stable cell line suitable for experimental applications. IB3-1 cells grow as an adherent monolayer, display an elongated morphology, and represent a well-established model for functional studies on the *CFTR* (*Cystic Fibrosis Transmembrane Conductance Regulator*) gene. Reverse-transcription (RT) PCR and Western blotting data showing high-level production of Spike mRNA and S-protein by BNT162b2-exposed IB3-1 cells can be found in the study published by Papi et al. (15). Effects of AGE, SAC and S1PC on pro-inflammatory genes induced by the BNT162b2 vaccine have been reported by Gasparello et al (14) and Papi et al (15).

The main purpose of this study was to determine (a) whether treatment of the IB3-1 bronchial epithelia cells with the BNT162b2 vaccine is associated with activation of the apoptotic pathway and (b) whether AGE, SAC and S1PC have any effect of therapeutic relevance on this experimental model system. In this study apoptosis was analyzed using the Annex V assay (measuring the level of phosphatidylserine flipper from the inner to the outer cell membrane leaflet) and the Caspase 3/7 assay. Notably, during apoptosis, the executioner Caspase-3 and Caspase-7 are activated and play a key role in flipping phosphatidylserine (PS) from the inner to the outer cell membrane. Externalized PS binds to specialized receptors and is responsible for the so-called “eat-me” signal for clearance by the immune system (22–24).

## RESULTS

### The experimental model system

In order to verify whether treatment of IB3-1 cells with AGE, SAC and S1PC alters the BNT162b2-induced apoptosis, cells have been treated with the BNT162b2 vaccine in the absence or in the presence of AGE, SAC and S1PC, as outlined in the pictorial representation shown in Fig. 1A. In the present study, we have exposed IB3-1 cells to BNT162b2 (0.7, 1.25, 2.5 and 3.5 mg/ml) and after 3 days of culture apoptosis was assessed using the Annexin V and Caspase 3/7 apoptosis assays (25,26) (Fig. 1B).

**Figure 1.**
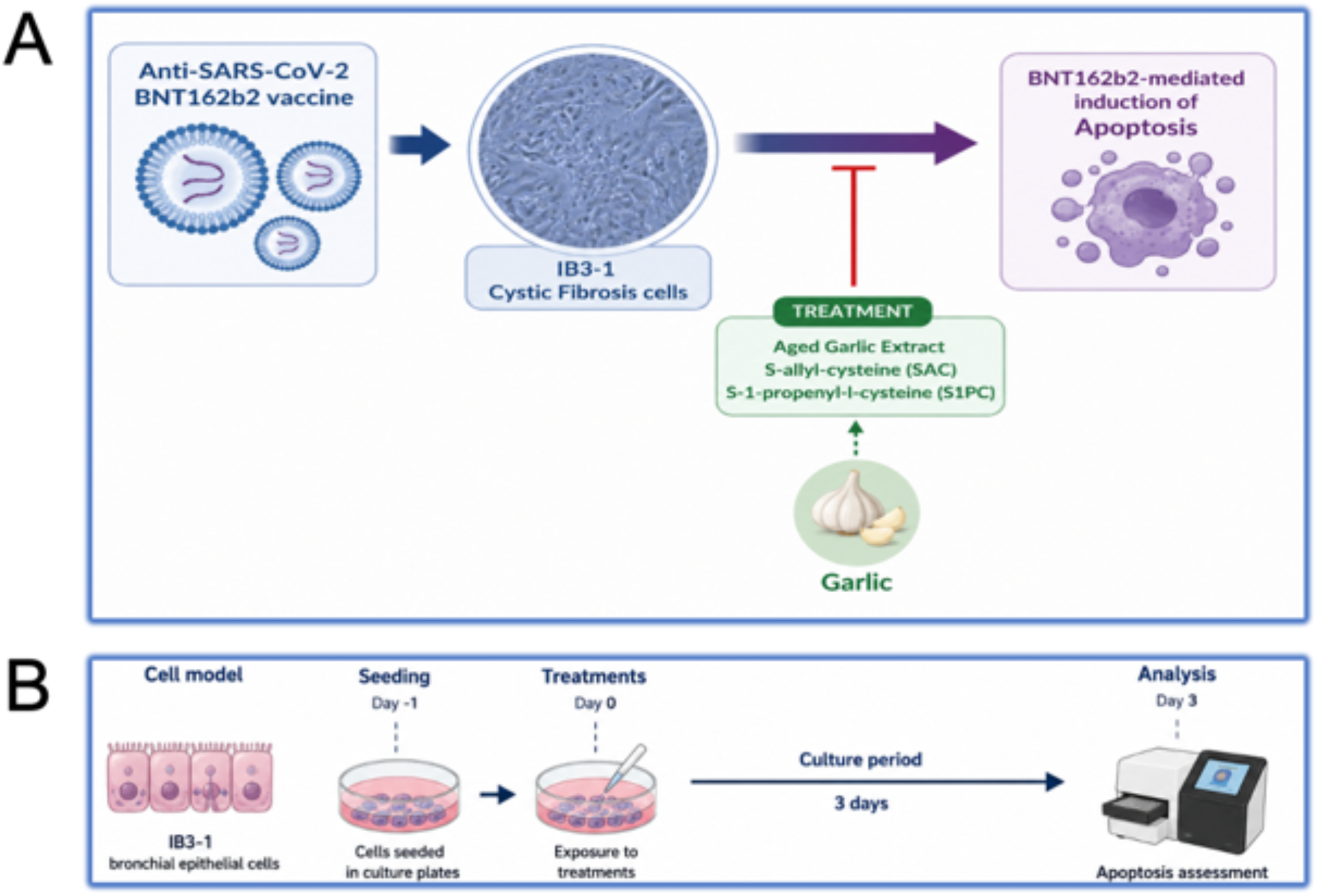
The experimental model system (A) and the employed protocol (B).

### Treatment of IB3-1 cells with the BNT162b2 vaccine is associated with high level of apoptosis

We have exposed IB3-1 cells (2x10^5^ cells/ml) to increasing amounts of BNT162b2 vaccine (0.7, 1.25, 2.5 and 3,5 μg/ml) and after 3 days of culture apoptosis was assessed using the Annex V apoptosis kit. Representative results are shown in Fig. 2A, outlining the fact that the proportion of late apoptotic/dead cells increased from 14.57% in untreated cells to 33.02% in IB3-1 cells treated with 1.25 μg/ml BNT162b2 and to 42.84% in cells treated with 3.5 μg/ml BNT162b2 (Fig. 2A). This preliminary experiment confirms previously published studies showing that SARS-CoV-2 Spike-producing vaccines induce apoptosis in different cellular systems (5–8). In Fig. 2 (panels B and C) the results from four independent experiments are shown, indicating that the decrease of live cells (Fig. 2B) and the increase of late apoptotic/dead cells (Fig. 2C) are highly significant (p < 0.001) after treatment of IB3-1 cells with 0.7 μg/ml of the BNT162b2 vaccine. A further, significant decrease of the proportion of live cells (Fig. 2B) and increase of the proportion of late apoptotic cells (Fig. 2C) were observed in cells treated with 1.25 μg/ml of the BNT162b2 vaccine. Notably, the BNT162b2 induced apoptosis of IB3-1 cells was confirmed using the Caspase 3/7 assay, as reported in Fig. 3. In the representative example shown in Fig. 3A, the proportion of late apoptotic/dead cells increased from 12.55% in untreated cells to 49.07% in IB3-1 cells treated with 1.25 μg/ml BNT162b2 and to 62.72% in cells treated with 3.5 μg/ml BNT162b2. In Fig. 3 (B and C) the results from four independent experiments are shown, indicating that the decrease of live cells (Fig. 3B) and the increase of late apoptotic/dead cells (Fig. 3C) are highly significant (p < 0.001) after treatment of IB3-1 cells with 0.7 μg/ml of the BNT162b2 vaccine. Also in this case, in agreement with the results of Fig. 2, a further, decrease of the percentage of live cells (Fig. 3B) and increase of the percentage of late apoptotic/dead cells (Fig. 3C) were observed in cells treated with 1.25 μg/ml of the BNT162b2 vaccine.

**Figure 2.**
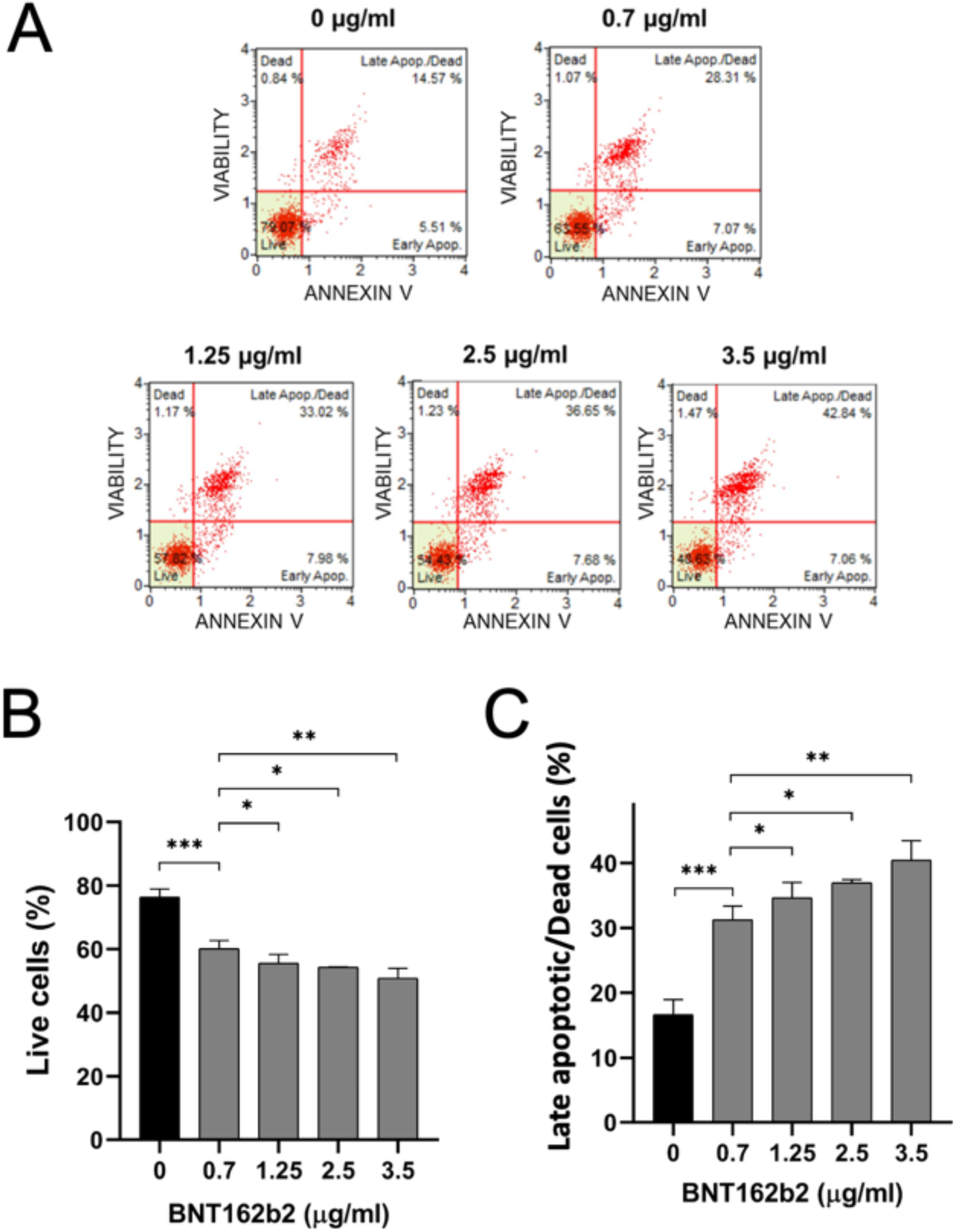
Treatment of IB3-1 cells with the BNT162b2 vaccine induces apoptosis: Annex V assay. A. Representative results on the effects of treatment of IB3-1 cells for 3 days in the presence of the indicated concentrations of BNT162b2 vaccine. Effects on apoptosis was analyzed by the Annexin V assay. The percentages of live, early apoptotic, late apoptotic and death cells are indicated on the plots. B, C. Summary of the effects of treatment of IB3-1 cells for 3 days in the presence of the indicated concentrations of the BNT162b2 vaccine. Live cells (B) and late apoptotic/dead cells (C) are indicated. Results represent the mean ± SD of four independent experiments. * = p < 0.05 (significant); ** = p < 0.01 (highly significant); *** = p < 0.001 (highly significant).

**Figure 3.**
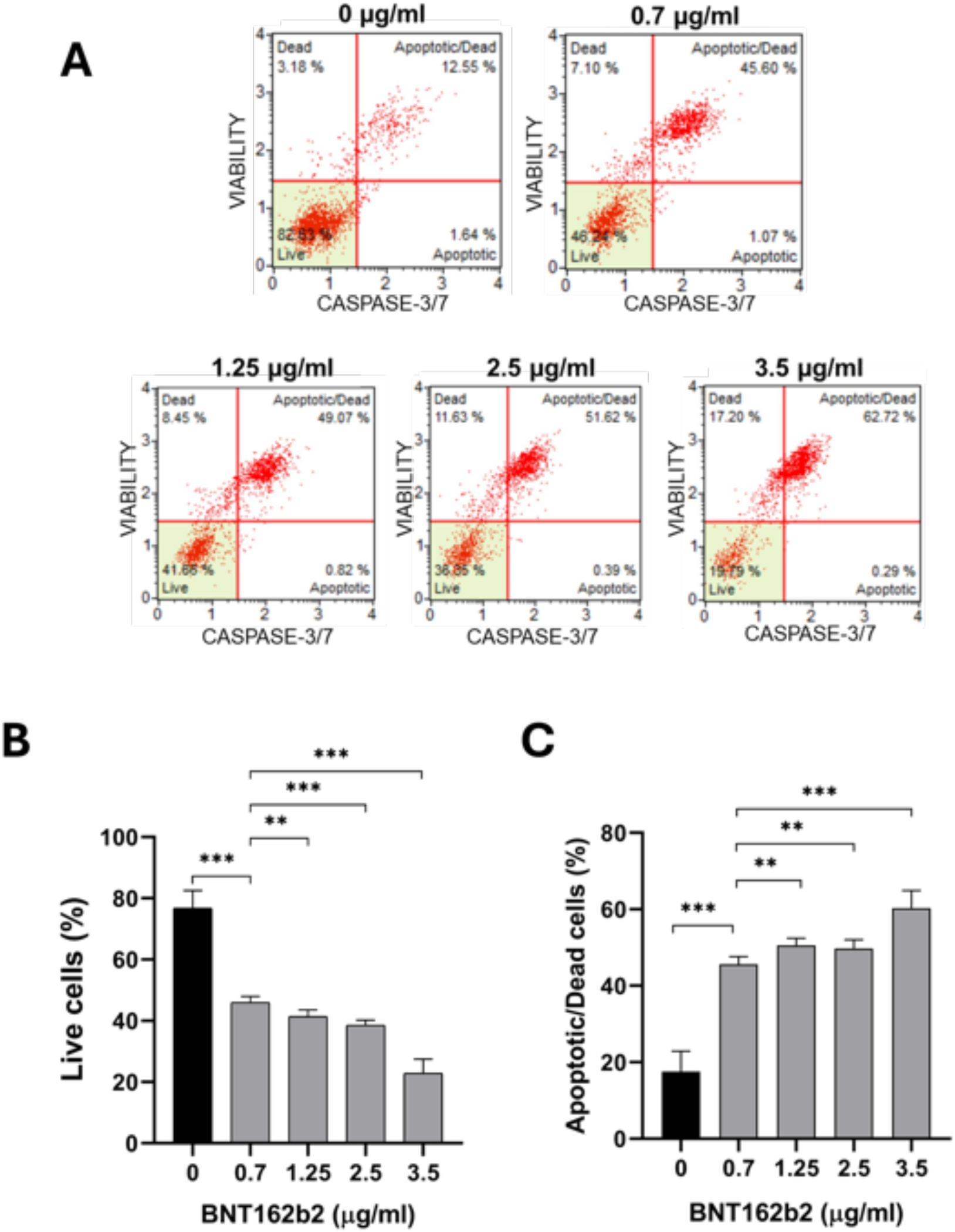
Treatment of IB3-1 cells with the BNT162b2 vaccine induces apoptosis: Caspase 3/7 assay. A. Representative results on the effects of treatment of IB3-1 cells for 3 days in the presence of the indicated concentrations of BNT162b2 vaccine. Apoptosis was assessed using the Caspase 3/7 assay. The percentages of live, early apoptotic, late apoptotic and death cells are indicated on the plots. B,C. Summary of the effects of treatment of IB3-1 cells for 3 days in the presence of the indicated concentrations of the BNT162b2 vaccine. Live cells (B) and late apoptotic/dead cells (C) are indicated. Results represents the mean ± SD of four independent experiments. * = p < 0.05 (significant); ** = p < 0.01 (highly significant); *** = p < 0.001 (highly significant).

Altogether, these results indicate that 0.7 and 1.25 μg/ml of the BNT162b2 vaccine are experimental conditions suitable to assess the effects of AGE, SAC and S1PC on BNT162b2-induced apoptosis.

### Treatment of IB3-1 cells with AGE, SAC and S1PC does not affect apoptosis

The purpose of the experiment shown in Fig. 4A was to verify whether treatment of IB3-1 cells with AGE, SAC and S1PC is associated with changes in the apoptosis profile. No significant changes in total apoptotic cells were observed in IB3-1 cells treated for 3 days with AGE, SAC and S1PC. Similar conclusion can be gathered when early apoptotic and late apoptotic cells were considered (data not shown). In parallel, we have performed MTT assays that excluded the presence of major cytotoxicity under the experimental conditions considered in this study, as shown in Fig.4B. No significant toxicity was found in IB3-1 cells treated for 3 days with 0.5 and 2 mg/ml AGE, 50 and 100 μM SAC and 50 and 100 μM S1PC. Decrease of viability was observed only at higher concentrations of the products (16 mg/ml AGE, 2 mM SAC and 2 mM S1PC (data not shown). It is worth noting that the experimental data obtained support the concept that the possible effects of AGE, SAC and S1PC did not represent a “confounding factor” when the effects of these agents were analyzed on BNT162b2-treated IB3-1 cells.

**Figure 4.**
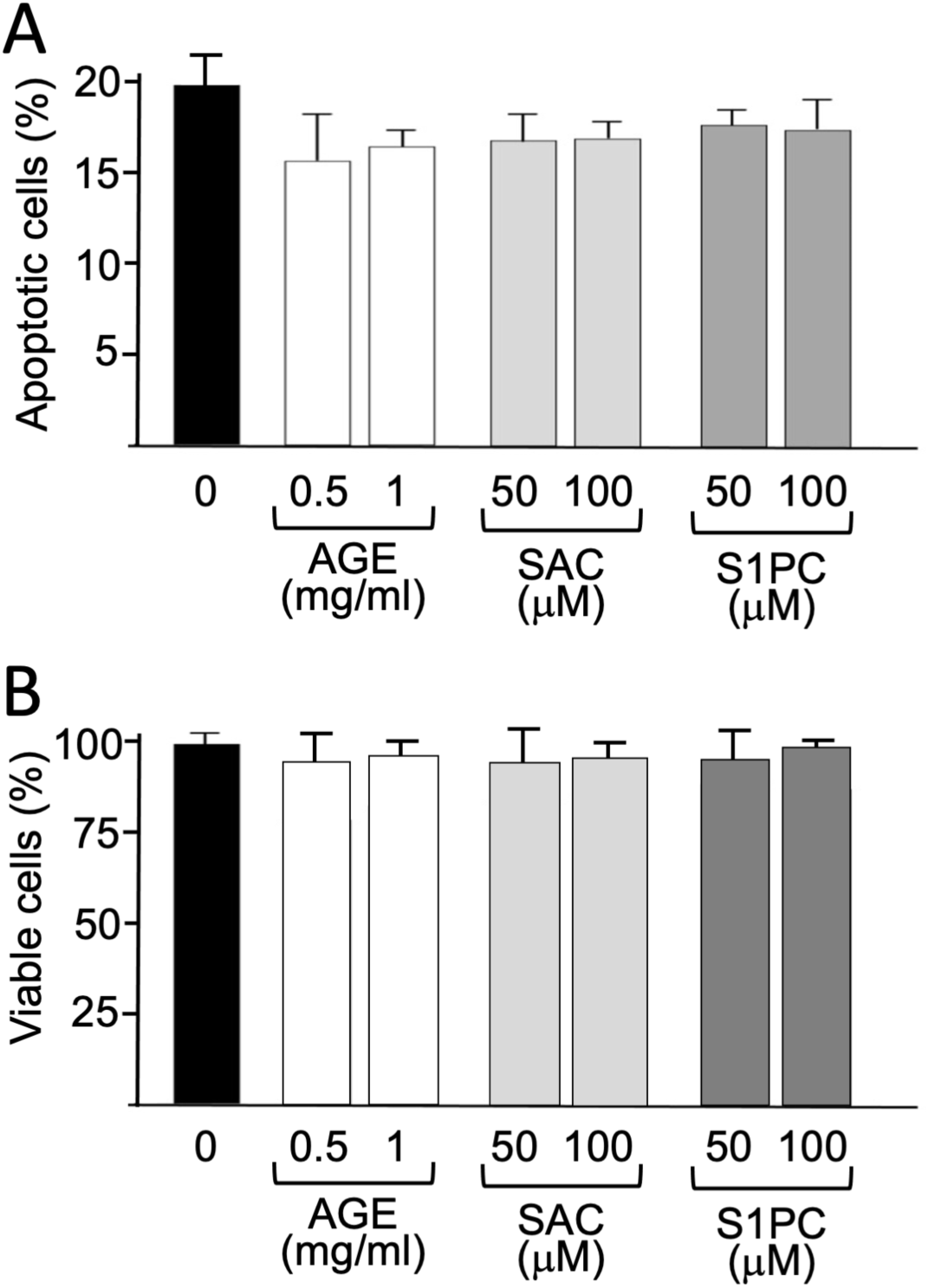
Effects of treatment of IB3-1 cells for 3 days in the presence of the indicated concentrations of AGE, SAC and S1PC. A. Effects on apoptosis (Annexin V assay). Changes in the percentage of apoptotic IB3-1 cells were found not significant (in all cases p > 0.05). B. Effects on viability (MTT assay). Changes in the percentage of viable IB3-1 cells were found not significant (in all cases p > 0.05). The results shown in panels A and B represent the average ± SD of three independent experiments.

### Treatment with AGE, SAC and S1PC partially reverses the BNT-162b2-induced apoptosis of IB3-1 cells

In order to verify whether treatment of IB3-1 cells with AGE, SAC and S1PC alters the BNT162b2-induced apoptosis of IB3-1 cells, cells have been treated with 1.25 μg/ml of the BNT162b2 vaccine and cultured for 3 days in the presence of 0.5 and 1 mg/ml of AGE, 50 and 100 μM of both SAC and S1PC. The representative experiment shown in Fig. 5 reports the results obtained using cells treated with 1.25 μg/ml of the BNT162b2 vaccine (see Fig. 5A) and cultured the presence of 1 mg/ml AGE (Fig. 5B) and 100 μM SAC (Fig. 5C) and S1PC (Fig. 5D). In this representative experiment the Annexin V assay was employed to assess apoptosis. Notably, the percentage of the cells in both the early and late phases of apoptosis is reduced in cells treated with the BNT162b2 vaccine in the presence of AGE, SAC and S1PC. In the case of late apoptotic/dead cells, the reduction was from 38.16% (IB3-1 cells treated with only the BNT162b2 vaccine) to 28.23% (BNT162b2 vaccine + AGE), 27.20% (BNT162b2 vaccine + SAC) and 27.42% (BNT162b2 vaccine + S1PC). Statistical analysis shows that the decrease of apoptotic cells is significant in four independent experiments carried out with IB3-1 cells treated with 1.25 μg/ml BNT162b2 vaccine (Fig. 5E).

**Figure 5.**
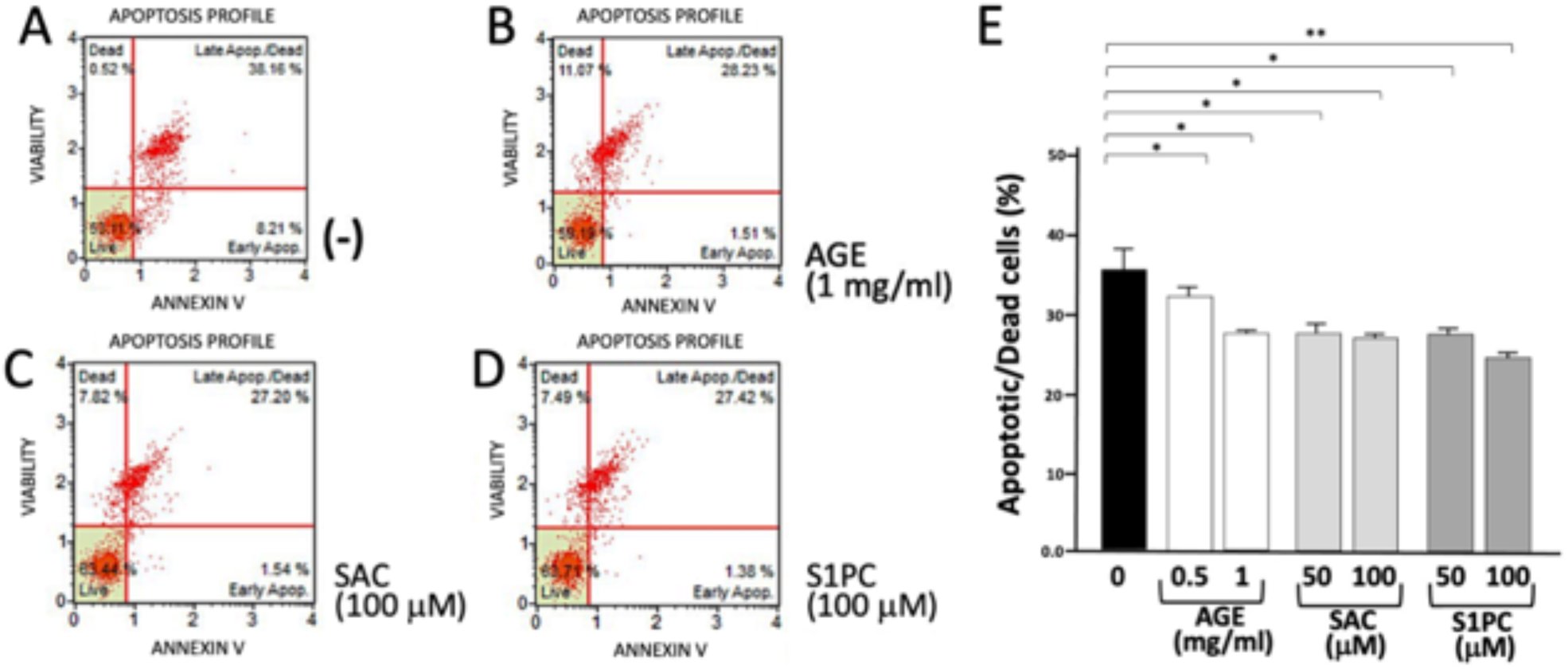
Impact of AGE, SAC, and S1PC on apoptosis in IB3-1 cells exposed to 1.25 μg/ml BNT162b2 vaccine: Annexin V assay. IB3-1 cells were treated for 72 hours with 1.25 μg/ml BNT162b2 in the presence of AGE (0.5 and 1 mg/ml), SAC (50 and 100 μM), and S1PC (50 and 100 μM), as specified. Apoptotic responses were assessed using the Annexin V assay. A-D. Representative results. E. Summary of the results obtained. Data are expressed as mean ± SD from four independent experiments. * = p < 0.05; ** = p < 0.01.

The effects of the AGE components SAC and S1PC in reducing the BNT162b2 induced apoptosis of IB3-1 cells was also confirmed using the Caspase 3/7 kit to assess apoptosis, as shown in Fig. 6. Using this analytical approach, the proportion of the cells in both the early and late phases of apoptosis was also reduced in cells treated with the BNT162b2 vaccine in the presence of AGE, SAC and S1PC. In the case of late apoptotic/dead cells, the reduction was from 52.60% (IB3-1 cells treated with only the BNT162b2 vaccine) to 46.36% (BNT162b2 vaccine + 100 μM SAC) and 42.03% (BNT162b2 vaccine + 100 μM S1PC). Statistical analysis shows that the decrease of apoptotic cells mediated by SAC and S1PC is significant in four independent experiments carried out with IB3-1 cells treated with 1.25 μg/ml BNT162b2 vaccine (Fig. 6E).

**Figure 6.**
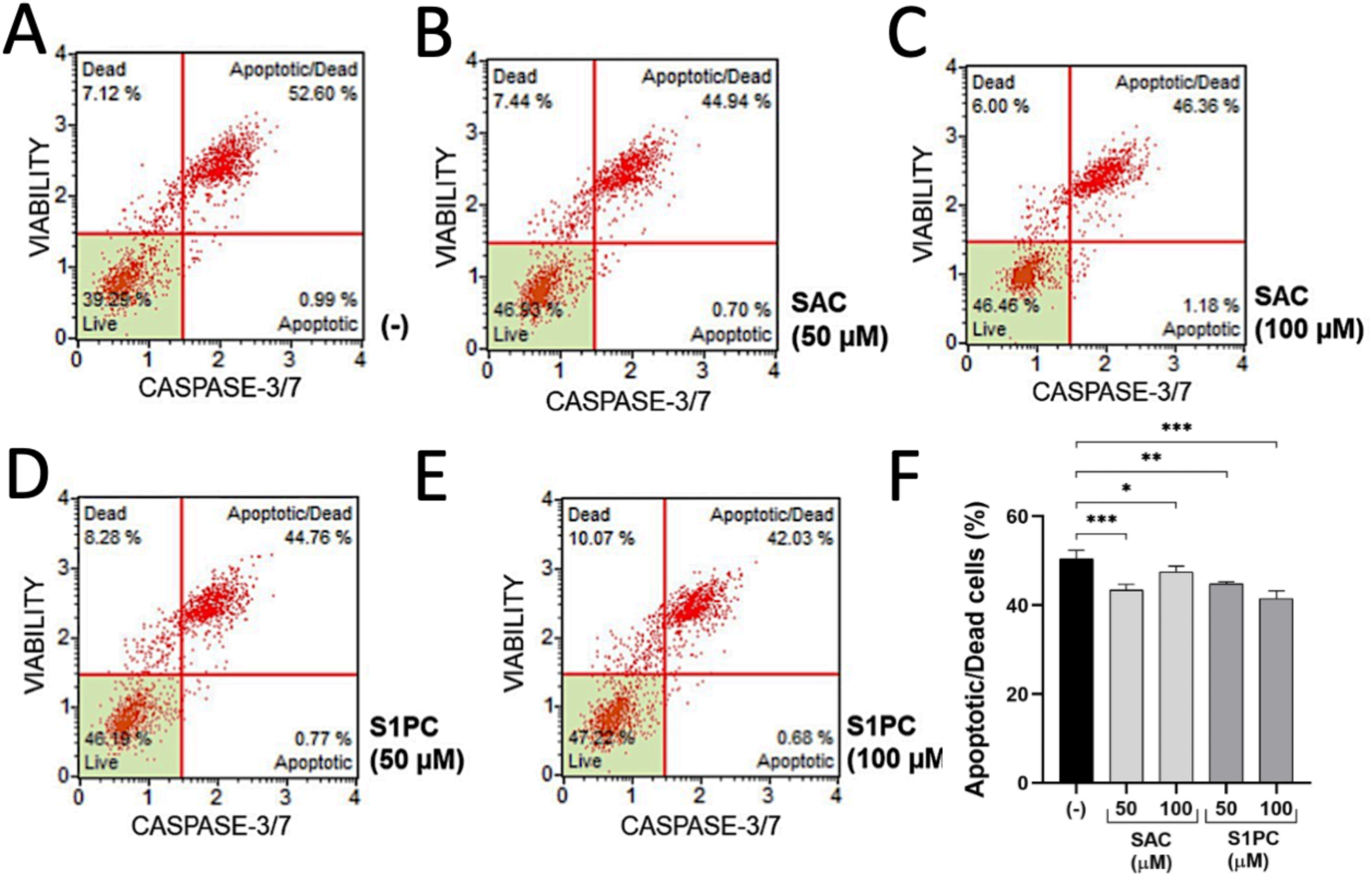
Impact of SAC, and S1PC on apoptosis in IB3-1 cells exposed to 1.25 μg/ml BNT162b2 vaccine: Caspase 3/7 assay. IB3-1 cells were treated for 72 hours with 1.25 μg/ml BNT162b2 in the presence of SAC (50 and 100 μM), and S1PC (50 and 100 μM), as specified. Apoptotic responses were assessed using the Caspase 3/7 assay. A-D. Representative results. F. Summary of the results obtained. Data are expressed as mean ± SD from four independent experiments. * = p < 0.05; ** = p < 0.01.

## DISCUSSION

One of the most powerful strategies employed against the COVID-19 outbreak is based on the use of anti-SARS-CoV-2 vaccines. Among the most used worldwide, the RNA-based vaccines Pfizer/BioNTech BNT162b2 (9) and Moderna mRNA-1273 (10) were demonstrated to be very effective and safe. These COVID-19 vaccines contain non-replicative forms of SARS-CoV-2 Spike mRNA, produce *in vitro* and *in vivo* large amounts of S-protein and stimulate high production of high levels of anti-SARS-CoV-2 antibodies. Review articles on the global pharmacological scientific efforts in the field of COVID-19 vaccine development are available (27–31), confirming that, thanks to the extensive use of COVID-19 vaccines and the improvement of the management of COVID-19 patients, the pandemic is at present under control. However, in addition to the beneficial effects of the Spike-mRNA based COVID-19 vaccines, side adverse effects were reported, strongly suggesting the need for further and more extensive evaluations of the short- and long-term effects of the COVID-19 vaccines on human tissue systems (1–4).

In this respect, the adverse effects of COVID-19 mRNA vaccines could be due to the presence of circulating spike, as discussed in several studies, such as those published by Trougakos et al. (1) and by Gopalaswamy et al. (3), who concluded that the potential ambivalence of human response post-COVID-19 vaccination necessitates mitigating the adverse side effects (3). In this respect, Meo et al. compared the biological, pharmacological characteristics and adverse effects of Pfizer/BioNTech and Moderna Vaccines. The expected conclusion is that the induction of anti SARS-CoV2 antibodies is associated with the production of high levels of the Spike protein (S-protein) (4).

One of the biological adverse effects of Spike protein (and, indirectly, of the Spike-producing vaccines) is high induction of apoptosis in different cellular systems (5–9). Notably, the induction of apoptosis by SARS-CoV-2 Spike protein is usually associated with deep alteration(s) of the cellular functions (9–12). With respect to this issue, Cesar et al. found that T4 apoptosis in the acute phase of SARS-CoV-2 infection predicts long COVID (32). Therefore, inhibiting apoptosis might be a strategy to mitigation some of the adverse side effects of Spike-producing RNA vaccines. A study sustaining this concept has been published by Roy et al. (13), who reported that *Eupatorium perfoliatum* prevents and alleviates SARS-CoV-2 Spike Protein-Induced lung inflammatory response and apoptosis (13).

The main result of our study is that Aged Garlic Extracts (AGE), and its components S-allyl-Cysteine (SAC) and S1-propenyl-l-Cysteine (S1PC^TM^) are able to mitigate the pro-apoptotic effects of the BNT162b2 vaccine and partially revert induction of early and late apoptotic cells in a bronchial cell line (the IB3-1 cell line) pretreated with the BNT162b2 vaccine. The efficacy of the reversion of BNT162b2 induced apoptosis was comparable to that found by Roy et al. using *Eupatorium perfoliatum* preparations. We therefore suggest that the natural extract AGE and its derivatives, SAC and S1PC, should be considered for drafting experimental protocols for counteracting apoptosis induced by the Spike mRNA-based COVID-19 vaccines.

Our study should be considered a pilot investigation, and the inhibitory effects of AGE, SAC and S1PC on BNT-162b2 induced apoptosis should be further studied by performing in-depth analysis of the expression of pro-apoptotic (Bax and Caspases-3 and 9) (33,34) and anti-apoptotic (Bcl-2) (35) genes. In order to fully understand the potential clinical applications of our study, the effects of AGE, SAC and S1PC should be also analyzed in other cellular systems exposed to the BNT162b2 vaccine, including cells present within the vaccination site (skeletal muscle cells, histiocytic cells and fibrocytes) (36) and B-lymphocytes (37) and T-lymphoid cells (38).

In addition, the possible effects of AGE, SAC and S1PC in mitigating induced apoptosis deserve to be investigated in other pathologies in which apoptosis should be counteracted. For instance, *Pseudomonas aeruginosa* infection triggers programmed cell death (including apoptosis, pyroptosis, necroptosis, ferroptosis, and PANoptosis) as demonstrated by the studies published by Phuong et al. (39) and Tan et al. (40).

Collectively, our study indicates that Aged Garlic Extracts (AGE), and its components S-allyl-Cysteine (SAC) and S1-propenyl-l-Cysteine (S1PC), are able to mitigate the pro-apoptotic effects of the BNT162b2 vaccine and partially revert induction of early and late apoptotic cells in a bronchial cell line (the IB3-1 cell line) pretreated with the BNT162b2 vaccine. The results of his study are fully in agreement with reports underlining the effects of garlic compounds on induced apoptosis in several cellular model systems (41–50).

The efficacy of the reversion of BNT162b2 induced apoptosis suggests that AGE, SAC and S1PC should be considered for the development of protocols for counteracting apoptosis induced by the Spike mRNA-based COVID-19 vaccines. This might be of potential clinical relevance, since one of the biological adverse effects of Spike protein (and, indirectly, of the Spike-producing vaccines) is high induction of apoptosis in different cellular systems.

## MATERIALS AND METHODS

### Materials

All reagents and chemicals were analytical grade. BNT162b2 (COMIRNATYTM, Lot. FP8191) was obtained from the Hospital Pharmacy of University of Padova (Roma, Italy).

### Extraction and Chemical Characterization of AGE, SAC and S1PC

AGE, SAC and S1PC^TM^ were provided by Wakunaga Pharmaceutical Co., Ltd. (Hiroshima, Japan) and manufactured as described by Kanamori et al. (51). Briefly, AGE was prepared by rinsing garlic (*Allium sativum* L.) cloves with purified water, slicing them and then soaking them in a water–ethanol mixture, which was then naturally extracted/aged for more than 10 months at room temperature. The AGE powder used in our experiments was prepared by lyophilization. It contained approximately 28.6% (w/v, 286 mg/ml) solid material, 0.63% (6.3 mg/ml) arginine and 0.1% SAC (calculated on a dry-weight basis) as a marker compound for standardization (52). AGE, SAC and S1PC powders were freshly dissolved in complete cell growth medium prior to each experiment. The chemical characterization of AGE, SAC and S1PC has been published elsewhere (14,15).

### Cell Culture Conditions

The human bronchial epithelial IB3-1 cell line was obtained from Thermo Fisher Scientific, Inc. and cultured in LHC-8 medium (Gibco, Thermo Fisher Scientific, Inc.; lot no. 3064217), specifically formulated for this cell line, supplemented with 5% fetal bovine serum (FBS; Biowest, Nuaillé, France), without antibiotics. Cultures were maintained in 25-cm^2^ tissue culture flasks or, when required for cell expansion, in 75-cm^2^ flasks, at 37°C in a humidified atmosphere containing 5% CO_2_ (53).

### Treatment of IB3-1 Cells with the BNT162b2 Vaccine and AGE, SAC or S1PC

For treatment with the BNT162b2 vaccine, IB3-1 cells were seeded at 50% of confluence and then treated with 1 μg/mL of the vaccine, as elsewhere described (54,55). After 24 h, cells were additionally treated with AGE, SAC and S1PC.

### Analysis of apoptosis: Annexin V/7-AAD apoptosis assay

Apoptosis of IB3-1 cells exposed to different concentrations of BNT162b2 and treated with the selected nutraceutical compounds was assessed using the Muse® Annexin V & Dead Cell Kit and the Muse® Cell Analyzer (Merck Millipore), according to the manufacturer’s instructions. Following treatment cells were detached and counted. For negative controls, 100 µl of cell suspension was mixed with 100 µl of Muse® Annexin V & Dead Cell Reagent. For treated samples, 50 µl of cell suspension was mixed with 50 µl of reagent. Samples were gently mixed and incubated for 20 min at room temperature in the dark. Samples were analyzed using the “Annexin V & Dead Cell” protocol, acquiring 2,000 events per sample. At least two consecutive acquisitions were performed for each sample. Muse® Analysis software was used to quantify apoptotic cell populations (56).

### Analysis of apoptosis: Caspase-3/7 activity assay

Caspase-3/7 activation was assessed in IB3-1 cells to validate the Annexin V assay results. The analysis was performed using the Muse® Caspase-3/7 Assay Kit and Muse® Cell Analyzer (Merck Millipore), according to the manufacturer’s instructions. Cells were detached using 100 µl of trypsin per well for 7 min, after which the enzyme was neutralized with 100 µl of FBS. DPBS (800 µl) was added to obtain a final volume of 1 ml, and the cell concentration was adjusted in order to achieve the range of 1x10^5^ - 6x10^6^ cells/ml. Samples were analyzed using the “Caspase-3/7” protocol. Samples were centrifuged at 1,200 rpm for 8 min at 4°C, and the resulting pellets were resuspended in 50 µl of 1X Assay Buffer. Five microlitres of Muse® Caspase-3/7 reagent, diluted 1:8 in 1X DPBS, was added to each sample. After incubation for 30 min at 37°C, 150 µl of Muse® 7-AAD working solution was added, followed by incubation for 5 min at room temperature in the dark. Samples were analyzed using the Muse® Cell Analyzer “Caspase-3/7” Software Modules (Millipore) (56).

### MTT cell viability assay

The cytotoxicity of AGE, SAC and S1PC compounds was assessed in IB3-1 cells using the MTT assay. Cells were seeded in 96-well plates at a density of 5 x 10^3^ cells/well in 100 µl of LHC-8 medium supplemented with 5% FBS. After cell adhesion, cells were treated with compounds at concentrations of 0.5 and 1 mg/ml AGE, 50 and 100 µM SAC, and 50 and 100 µM S1PC. Untreated cells, vehicle-treated cells, and H_2_O_2_ treated (0.1% final concentration) cells were included as controls. After 72 h of treatment, 20 µl of MTT solution (5 mg/ml in sterile 1X PBS; Merck, cat. no. 1003688598) was added directly to each well. Plates were incubated for 3.5 h at 37°C in the dark. The medium was then removed, and the resulting formazan crystals were dissolved in 200 µl of DMSO by shaking the plates for 15 min in the dark. Absorbance was measured at 570 nm, using 650 nm as the reference wavelength, with a Tecan Spark^®^ microplate reader (Tecan Group Ltd.). Background-corrected absorbance values were calculated by subtracting the absorbance at 650 nm from that measured at 570 nm (A_570 −_ A_650_). Cell viability was expressed as a percentage relative to untreated control cells (57).

### Statistical analysis

The data are presented as mean ± SD of at least three independent experiments. Statistical differences between groups were analyzed using one-way ANOVA. Prism (v. 8.02) by GraphPad software (Dotmatics) was used (followed by Bonferroni’s test). P<0.05 was considered to indicate a significant difference.

## ACKNOWLEDGEMENTS

This work was funded by by Wakunaga Pharmaceutical Co. Ltd. (Hiroshima, Japan) (to E.A., A.F. and R.G.). AF and RG were also funded by the MUR-FISR COVID-miRNAPNA Project (FISR2020IP_04128) (to R.G. and A.F.), by CIB (grant CIB-2024-AF) and by FIRD 2024 funds from the University of Ferrara to A.F. (grant n° FIRD-Finotti 2024). A.F. received funds from the Interuniversity Consortium for Biotechnologies, Italy (CIB) (grant no. CIB-MUR-Unife2024). The authors would like to thank Dr Takahiro Ogawa and Dr Toshiaki Matsutomo (Wakunaga Pharmaceutical Co., Ltd.) for organizing the shipment of AGE, SAC and S1PC. We thank the International Polyamine Foundation for the administrative management of the Project.

## REFERENCES

1. Trougakos IP, Terpos E, Alexopoulos H, Politou M, Paraskevis D, Scorilas A, Kastritis E, Andreakos E, Dimopoulos MA. Adverse effects of COVID-19 mRNA vaccines: the spike hypothesis. Trends Mol Med. 2022 Jul;28(7):542–554. doi: 10.1016/j.molmed.2022.04.007. Epub 2022 Apr 21.

2. Cosentino M, Marino F. Understanding the Pharmacology of COVID-19 mRNA Vaccines: Playing Dice with the Spike? Int J Mol Sci. 2022 Sep 17;23(18):10881. doi: 10.3390/ijms231810881.

3. Gopalaswamy R, Aravindhan V, Subbian S. The Ambivalence of Post COVID-19 Vaccination Responses in Humans. Biomolecules. 2024 Oct 17;14(10):1320. doi: 10.3390/biom14101320.

4. Meo SA, Bukhari IA, Akram J, Meo AS, Klonoff DC. COVID-19 vaccines: comparison of biological, pharmacological characteristics and adverse effects of Pfizer/BioNTech and Moderna Vaccines. Eur Rev Med Pharmacol Sci. 2021 Feb;25(3):1663–1669. doi: 10.26355/eurrev_202102_24877.

5. Li F, Li J, Wang PH, Yang N, Huang J, Ou J, Xu T, Zhao X, Liu T, Huang X, Wang Q, Li M, Yang L, Lin Y, Cai Y, Chen H, Zhang Q. SARS-CoV-2 spike promotes inflammation and apoptosis through autophagy by ROS-suppressed PI3K/AKT/mTOR signaling. Biochim Biophys Acta Mol Basis Dis. 2021 Dec 1;1867(12):166260. doi: 10.1016/j.bbadis.2021.166260. Epub 2021 Aug 27.

6. Barhoumi T, Alghanem B, Shaibah H, Mansour FA, Alamri HS, Akiel MA, Alroqi F, Boudjelal M. SARS-CoV-2 Coronavirus Spike Protein-Induced Apoptosis, Inflammatory, and Oxidative Stress Responses in THP-1-Like-Macrophages: Potential Role of Angiotensin-Converting Enzyme Inhibitor (Perindopril). Front Immunol. 2021 Sep 20;12:728896. doi: 10.3389/fimmu.2021.728896. eCollection 2021.

7. Yamamoto T, Koyama Y, Ujita T, Sawada E, Kishimoto N, Seo K. SARS-CoV-2 recombinant spike protein induces cell apoptosis in rat taste buds. J Dent Sci. 2023 Jan;18(1):428–431. doi: 10.1016/j.jds.2022.08.016. Epub 2022 Aug 26.

8. Wu CT, Lidsky PV, Xiao Y, Lee IT, Cheng R, Nakayama T, Jiang S, Demeter J, Bevacqua RJ, Chang CA, Whitener RL, Stalder AK, Zhu B, Chen H, Goltsev Y, Tzankov A, Nayak JV, Nolan GP, Matter MS, Andino R, Jackson PK. SARS-CoV-2 infects human pancreatic β cells and elicits β cell impairment. Cell Metab. 2021 Aug 3;33(8):1565–1576.e5. doi: 10.1016/j.cmet.2021.05.013. Epub 2021 May 18.

9. Polack FP, Thomas SJ, Kitchin N, Absalon J, Gurtman A, Lockhart S, Perez JL, Pérez Marc G, Moreira ED, Zerbini C, Bailey R, Swanson KA, Roychoudhury S, Koury K, Li P, Kalina WV, Cooper D, Frenck RW Jr, Hammitt LL, Türeci Ö, Nell H, Schaefer A, Ünal S, Tresnan DB, Mather S, Dormitzer PR, Şahin U, Jansen KU, Gruber WC; C4591001 Clinical Trial Group. Safety and Efficacy of the BNT162b2 mRNA Covid-19 Vaccine. N Engl J Med. 2020 Dec 31;383(27):2603–2615. doi: 10.1056/NEJMoa2034577.

10. Baden LR, El Sahly HM, Essink B, Kotloff K, Frey S, Novak R, Diemert D, Spector SA, Rouphael N, Creech CB, McGettigan J, Khetan S, Segall N, Solis J, Brosz A, Fierro C, Schwartz H, Neuzil K, Corey L, Gilbert P, Janes H, Follmann D, Marovich M, Mascola J, Polakowski L, Ledgerwood J, Graham BS, Bennett H, Pajon R, Knightly C, Leav B, Deng W, Zhou H, Han S, Ivarsson M, Miller J, Zaks T; COVE Study Group. Efficacy and Safety of the mRNA-1273 SARS-CoV-2 Vaccine .N Engl J Med. 2021 Feb 4;384(5):403–416. doi: 10.1056/NEJMoa2035389. Epub 2020 Dec 30.

11. Gimenez S, Hamrouni E, André S, Picard M, Soundaramourty C, Lozano C, Vincent T, Tran TA, Kundura L, Estaquier J, Corbeau P. Monocytic reactive oxygen species-induced T-cell apoptosis impairs cellular immune response to SARS-CoV-2 mRNA vaccine. J Allergy Clin Immunol. 2025 Jan 10:S0091–6749(25)00011-9.

12. André S, Azarias da Silva M, Picard M, Alleaume-Buteau A, Kundura L, Cezar R, Soudaramourty C, André SC, Mendes-Frias A, Carvalho A, Capela C, Pedrosa J, Gil Castro A, Loubet P, Sotto A, Muller L, Lefrant JY, Roger C, Claret PG, Duvnjak S, Tran TA, Zghidi-Abouzid O, Nioche P, Silvestre R, Corbeau P, Mammano F, Estaquier J. Low quantity and quality of anti-spike humoral response is linked to CD4 T-cell apoptosis in COVID-19 patients. Cell Death Dis. 2022 Aug 27;13(8):741. doi: 10.1038/s41419-022-05190-0.

13. Roy A, Sarkar A, Roy AK, Ghorai T, Nayak D, Kaushik S, Das S. Ultradiluted Eupatorium perfoliatum Prevents and Alleviates SARS-CoV-2 Spike Protein-Induced Lung Pathogenesis by Regulating Inflammatory Response and Apoptosis. Diseases. 2025 Jan 30;13(2):36. doi: 10.3390/diseases13020036.

14. Gasparello J, Papi C, Marzaro G, Macone A, Zurlo M, Finotti A, Agostinelli E, Gambari R. Aged Garlic Extract (AGE) and Its Constituent S-Allyl-Cysteine (SAC) Inhibit the Expression of Pro-Inflammatory Genes Induced in Bronchial Epithelial IB3-1 Cells by Exposure to the SARS-CoV-2 Spike Protein and the BNT162b2 Vaccine. Molecules. 2024 Dec 16;29(24):5938. doi: 10.3390/molecules29245938.

15. Papi C, Gasparello J, Marzaro G, Macone A, Zurlo M, Di Padua F, Fino P, Agostinelli E, Gambari R, Finotti A. S-1-propenyl-l-cysteine (S1PC), a major constituent of Aged Garlic Extract (AGE), exhibits inhibitory effects on pro-inflammatory gene expression in bronchial epithelial IB3-1 cells exposed to the BNT162b2 vaccine. Exp Ther Med 2025, in the press.

16. Aboudounya MM, Heads RJ. COVID-19 and Toll-Like Receptor 4 (TLR4): SARS-CoV-2 May Bind and Activate TLR4 to Increase ACE2 Expression, Facilitating Entry and Causing Hyperinflammation. Mediators Inflamm. 2021 Jan 14;2021:8874339. doi: 10.1155/2021/8874339. eCollection 2021.

17. Sahanic S, Hilbe R, Dünser C, Tymoszuk P, Löffler-Ragg J, Rieder D, Trajanoski Z, Krogsdam A, Demetz E, Yurchenko M, Fischer C, Schirmer M, Theurl M, Lener D, Hirsch J, Holfeld J, Gollmann-Tepeköylü C, Zinner CP, Tzankov A, Zhang SY, Casanova JL, Posch W, Wilflingseder D, Weiss G, Tancevski I. SARS-CoV-2 activates the TLR4/MyD88 pathway in human macrophages: A possible correlation with strong pro-inflammatory responses in severe COVID-19. Heliyon. 2023 Nov 17;9(11):e21893. doi: 10.1016/j.heliyon.2023.e21893. eCollection 2023 Nov.

18. Zhao Y, Kuang M, Li J, Zhu L, Jia Z, Guo X, Hu Y, Kong J, Yin H, Wang X, You F. SARS-CoV-2 spike protein interacts with and activates TLR41. Cell Res. 2021 Jul;31(7):818–820. doi: 10.1038/s41422-021-00495-9. Epub 2021 Mar 19.

19. Gasparello J, D’Aversa E, Papi C, Gambari L, Grigolo B, Borgatti M, Finotti A, Gambari R. Sulforaphane inhibits the expression of interleukin-6 and interleukin-8 induced in bronchial epithelial IB3-1 cells by exposure to the SARS-CoV-2 Spike protein. Phytomedicine. 2021 Jul;87:153583. doi: 10.1016/j.phymed.2021.153583. Epub 2021 May 4.

20. 20. Gasparello J, d’Aversa E, Breveglieri G, Borgatti M, Finotti A, Gambari R. In vitro induction of interleukin-8 by SARS-CoV-2 Spike protein is inhibited in bronchial epithelial IB3-1 cells by a miR-93-5p agomiR. Int Immunopharmacol. 2021 Dec;101(Pt B):108201. doi: 10.1016/j.intimp.2021.108201. Epub 2021 Sep 28.

21. Reddel RR, Ke Y, Gerwin BI, McMenamin MG, Lechner JF, Su RT, Brash DE, Park JB, Rhim JS, Harris CC. Transformation of human bronchial epithelial cells by infection with SV40 or adenovirus-12 SV40 hybrid virus, or transfection via strontium phosphate coprecipitation with a plasmid containing SV40 early region genes. Cancer Res. 1988 Apr 1;48(7):1904–9.

22. Mariño G, Kroemer G. Mechanisms of apoptotic phosphatidylserine exposure. Cell Res. 2013 Nov;23(11):1247–8. doi: 10.1038/cr.2013.115. Epub 2013 Aug 27.

23. Subham Preetam, Arunima Pandey, Richa Mishra, Gautam Mohapatra, Pratyasa Rath, Sumira Malik, Sarvesh Rustagi, Alisha Dash, Shailesh Kumar Samal. Phosphatidylserine: paving the way for a new era in cancer therapies. Mater. Adv. 2024; 5 (21): 8384–8403. 10.1039/d4ma00511b

24. Segawa K, Kurata S, Yanagihashi Y, Brummelkamp TR, Matsuda F, Nagata S. Caspase-mediated cleavage of phospholipid flippase for apoptotic phosphatidylserine exposure. Science. 2014 Jun 6;344(6188):1164–8. doi: 10.1126/science.1252809.

25. Kari S, Subramanian K, Altomonte IA, Murugesan A, Yli-Harja O, Kandhavelu M. Programmed cell death detection methods: a systematic review and a categorical comparison. Apoptosis. 2022 Aug;27(7-8):482–508. doi: 10.1007/s10495-022-01735-y. Epub 2022 Jun 17.

26. Shim MK, Yoon HY, Lee S, Jo MK, Park J, Kim JH, Jeong SY, Kwon IC, Kim K. Caspase-3/-7-Specific Metabolic Precursor for Bioorthogonal Tracking of Tumor Apoptosis. Sci Rep. 2017 Nov 30;7(1):16635. doi: 10.1038/s41598-017-16653-2.

27. Azeem M, Cancemi P, Mukhtar F, Marino S, Peri E, Di Prima G, De Caro V. Efficacy and limitations of SARS-CoV-2 vaccines - A systematic review. Life Sci. 2025 Apr 4;371:123610. doi: 10.1016/j.lfs.2025.123610. Online ahead of print.

28. Wang X, Pahwa A, Bausch-Jurken MT, Chitkara A, Sharma P, Malmenäs M, Vats S, Whitfield MG, Lai KZH, Dasari P, Gupta R, Nassim M, Van de Velde N, Green N, Beck E. Comparative Effectiveness of mRNA-1273 and BNT162b2 COVID-19 Vaccines Among Adults with Underlying Medical Conditions: Systematic Literature Review and Pairwise Meta-Analysis Using GRADE. Adv Ther. 2025 May;42(5):2040–2077. doi: 10.1007/s12325-025-03117-7. Epub 2025 Mar 10.

29. Wong BK, Mabbott NA. Systematic review and meta-analysis of COVID-19 mRNA vaccine effectiveness against hospitalizations in adults. Immunother Adv. 2024 Nov 27;4(1):ltae011. doi: 10.1093/immadv/ltae011. eCollection 2024.

30. Wilburn J, Sappe B, Jorge K, Hickey L, Nandyala D, Chadha T. Effectiveness of Pfizer Vaccine BNT162b2 Against SARS-CoV-2 in Americans 16 and Older: A Systematic Review. Cureus. 2024 Jul 22;16(7):e65111. doi: 10.7759/cureus.65111. eCollection 2024 Jul.

31. Katoto PD, Tamuzi JL, Brand AS, Marangu DM, Byamungu LN, Wiysonge CS, Gray G. Effectiveness of COVID-19 Pfizer-BioNTech (BNT162b2) mRNA vaccination in adolescents aged 12-17 years: A systematic review and meta-analysis. Hum Vaccin Immunother. 2023 Dec 31;19(1):2214495. doi: 10.1080/21645515.2023.2214495. Epub 2023 Jun 5.

32. Cezar R, Kundura L, André S, Lozano C, Vincent T, Muller L, Lefrant JY, Roger C, Claret PG, Duvnjak S, Loubet P, Sotto A, Tran TA, Estaquier J, Corbeau P. T4 apoptosis in the acute phase of SARS-CoV-2 infection predicts long COVID. Front Immunol. 2024 Jan 3;14:1335352. doi: 10.3389/fimmu.2023.1335352. eCollection 2023.

33. Pawlowski J, Kraft AS. Bax-induced apoptotic cell death. Proc Natl Acad Sci U S A. 2000 Jan 18;97(2):529–31. doi: 10.1073/pnas.97.2.529.

34. Johnson CR, Jarvis WD. Caspase-9 regulation: an update. Apoptosis. 2004 Jul;9(4):423–7. doi: 10.1023/B:APPT.0000031457.90890.13.

35. Edlich F. BCL-2 proteins and apoptosis: Recent insights and unknowns. Biochem Biophys Res Commun. 2018 May 27;500(1):26–34. doi: 10.1016/j.bbrc.2017.06.190. Epub

36. Beck A, Dietenberger H, Kunz SN, Mellert K, Möller P. Emergence of SARS-CoV-2 spike protein at the vaccination site. Immun Inflamm Dis. 2023 Mar;11(3):e827. doi: 10.1002/iid3.827.

37. Chen S, Guan F, Candotti F, Benlagha K, Camara NOS, Herrada AA, James LK, Lei J, Miller H, Kubo M, Ning Q, Liu C. The role of B cells in COVID-19 infection and vaccination. Front Immunol. 2022 Aug 30;13:988536. doi: 10.3389/fimmu.2022.988536. eCollection 2022.

38. Wang L, Nicols A, Turtle L, Richter A, Duncan CJ, Dunachie SJ, Klenerman P, Payne RP. T cell immune memory after covid-19 and vaccination. BMJ Med. 2023 Nov 22;2(1):e000468. doi: 10.1136/bmjmed-2022-000468. eCollection 2023.

39. Phuong MS, Hernandez RE, Wolter DJ, Hoffman LR, Sad S. Impairment in inflammasome signaling by the chronic Pseudomonas aeruginosa isolates from cystic fibrosis patients results in an increase in inflammatory response. Cell Death Dis. 2021 Mar 4;12(3):241. doi: 10.1038/s41419-021-03526-w.

40. Tan C, Luo Y. Advances in Pseudomonas aeruginosa-Induced Programmed Cell Death and Potential Targeted Treatment Strategies. Microorganisms. 2025 Nov 10;13(11):2560. doi: 10.3390/microorganisms13112560.

41. Huang XP, Shi ZH, Ming GF, Xu DM, Cheng SQ. S-Allyl-L-cysteine (SAC) inhibits copper-induced apoptosis and cuproptosis to alleviate cardiomyocyte injury. Biochem Biophys Res Commun. 2024 Oct 20;730:150341. doi: 10.1016/j.bbrc.2024.150341

42. Sakayanathan P, Loganathan C, Thayumanavan P. Protection of pancreatic beta cells against high glucose-induced toxicity by astaxanthin-s-allyl cysteine diester: alteration of oxidative stress and apoptotic-related protein expression. Arch Physiol Biochem. 2024 Jun;130(3):316–324. doi: 10.1080/13813455.2022.2064878.

43. Shao Z, Pan Z, Lin J, Zhao Q, Wang Y, Ni L, Feng S, Tian N, Wu Y, Sun L, Gao W, Zhou Y, Zhang X, Wang X. S-allyl cysteine reduces osteoarthritis pathology in the tert-butyl hydroperoxide-treated chondrocytes and the destabilization of the medial meniscus model mice via the Nrf2 signaling pathway. Aging (Albany NY). 2020 Oct 7;12(19):19254–19272. doi: 10.18632/aging.103757.

44. Chen P, Chen C, Hu M, Cui R, Liu F, Yu H, Ren Y. S-allyl-L-cysteine protects hepatocytes from indomethacin-induced apoptosis by attenuating endoplasmic reticulum stress. FEBS Open Bio. 2020 Sep;10(9):1900–1911. doi: 10.1002/2211-5463.12945. Epub 2020 Aug 16.

45. Sun ZW, Chen C, Wang L, Li YD, Hu ZL. S-allyl cysteine protects retinal pigment epithelium cells from hydroquinone-induced apoptosis through mitigating cellular response to oxidative stress. Eur Rev Med Pharmacol Sci. 2020 Feb;24(4):2120–2128. doi: 10.26355/eurrev_202002_20392.

46. Chen P, Hu M, Liu F, Yu H, Chen C. S-allyl-l-cysteine (SAC) protects hepatocytes from alcohol-induced apoptosis. FEBS Open Bio. 2019 Jul;9(7):1327–1336. doi: 10.1002/2211-5463.12684. Epub 2019 Jun 17.

47. Basu C, Sur R. S-Allyl Cysteine Alleviates Hydrogen Peroxide Induced Oxidative Injury and Apoptosis through Upregulation of Akt/Nrf-2/HO-1 Signaling Pathway in HepG2 Cells. Biomed Res Int. 2018 Nov 1;2018:3169431. doi: 10.1155/2018/3169431. eCollection 2018.

48. Orozco-Ibarra M, Muñoz-Sánchez J, Zavala-Medina ME, Pineda B, Magaña-Maldonado R, Vázquez-Contreras E, Maldonado PD, Pedraza-Chaverri J, Chánez-Cárdenas ME. Aged garlic extract and S-allylcysteine prevent apoptotic cell death in a chemical hypoxia model. Biol Res. 2016 Feb 1;49:7. doi: 10.1186/s40659-016-0067-6.

49. Mandal S, Mukherjee S, Chowdhury KD, Sarkar A, Basu K, Paul S, Karmakar D, Chatterjee M, Biswas T, Sadhukhan GC, Sen G. S-allyl cysteine in combination with clotrimazole downregulates Fas induced apoptotic events in erythrocytes of mice exposed to lead. Biochim Biophys Acta. 2012 Jan;1820(1):9–23. doi: 10.1016/j.bbagen.2011.09.019.

50. Kalayarasan S, Sriram N, Sureshkumar A, Sudhandiran G. Chromium (VI)-induced oxidative stress and apoptosis is reduced by garlic and its derivative S-allylcysteine through the activation of Nrf2 in the hepatocytes of Wistar rats. J Appl Toxicol. 2008 Oct;28(7):908–19. doi: 10.1002/jat.1355.

51. Kanamori, Y.; Via, L.D.; Macone, A.; Canettieri, G.; Greco, A.; Toninello, A.; Agostinelli, E. Aged garlic extract and its constituent, S-allyl-L-cysteine, induce the apoptosis of neuroblastoma cancer cells due to mitochondrial membrane depolarization. Exp. Ther. Med. 2020, 19, 1511–1521.

52. Bar M, Binduga UE, Szychowski KA. Methods of Isolation of Active Substances from Garlic (Allium sativum L.) and Its Impact on the Composition and Biological Properties of Garlic Extracts. Antioxidants (Basel). 2022 Jul 9;11(7):1345. doi: 10.3390/antiox11071345.

53. Borgatti M, Mazzitelli S, Breveglieri G, Gambari R, Nastruzzi C. Induction by TNF-α of IL-6 and IL-8 in cystic fibrosis bronchial IB3-1 epithelial cells encapsulated in alginate microbeads. J Biomed Biotechnol. 2010;2010:907964. doi: 10.1155/2010/907964. Epub 2010 Sep 8.

54. Zurlo M, Gasparello J, Verona M, Papi C, Cosenza LC, Finotti A, Marzaro G, Gambari R. The anti-SARS-CoV-2 BNT162b2 vaccine suppresses mithramycin-induced erythroid differentiation and expression of embryo-fetal globin genes in human erythroleukemia K562 cells. Exp Cell Res. 2023 Dec 15;433(2):113853. doi: 10.1016/j.yexcr.2023.113853. Epub 2023 Nov 7.

55. Cosenza LC, Marzaro G, Zurlo M, Gasparello J, Zuccato C, Finotti A, Gambari R. Inhibitory effects of SARS-CoV-2 spike protein and BNT162b2 vaccine on erythropoietin-induced globin gene expression in erythroid precursor cells from patients with beta-thalassemia. Exp Hematol. 2024 Jan;129:104128. doi: 10.1016/j.exphem.2023.11.002. Epub 2023 Nov 7.

56. Gasparello J, Papi C, Zurlo M, Gambari L, Manicardi A, Rozzi A, Ferrarini M, Corradini R, Gambari R, Finotti A. MicroRNAs miR-584-5p and miR-425-3p Are Up-Regulated in Plasma of Colorectal Cancer (CRC) Patients: Targeting with Inhibitor Peptide Nucleic Acids Is Associated with Induction of Apoptosis in Colon Cancer Cell Lines. Cancers (Basel). 2022 Dec 25;15(1):128. doi: 10.3390/cancers15010128.

57. Gasparello J, Papi C, Zurlo M, Volpi S, Gambari R, Corradini R, Casnati A, Sansone F, Finotti A. Cationic Calix[4]arene Vectors to Efficiently Deliver AntimiRNA Peptide Nucleic Acids (PNAs) and miRNA Mimics. Pharmaceutics. 2023 Aug 10;15(8):2121. doi: 10.3390/pharmaceutics15082121.

